# Transposable element activity and polymorphisms drive structural variability within and between individual in bivalves

**DOI:** 10.64898/2026.02.10.705023

**Authors:** Jacopo Martelossi, Andrea Luchetti, Alexander Suh, Fabrizio Ghiselli, Valentina Peona

**Author notes:** corresponding author: Jacopo Martelossi.

## Abstract

Structural variants (SVs) represent one of the most abundant sources of genetic variation across eukaryotes, with transposable elements (TEs) standing out as primary contributors in their emergence. While the growing availability of chromosome-scale genomes has revealed the central role of SVs in species diversification and adaptation, non-model invertebrates remain critically understudied in this context. Here, we explore how SVs and their interaction with TEs shape genetic diversity at both the individual and population levels. To achieve this, we leverage four high-quality oyster genomes and a large-scale dataset of the Estuarine oyster (*Crassostrea ariakensis*) collected across a wide range of different temperature and salinity conditions. We explicitly account for strengths and limitations of current SV-calling software, and benchmark our results through simulations. We uncover pervasive within-individual structural variability, with up to 14% of oyster genome basepairs being in an hemizygous state. The strong enrichment of TEs within these SVs is driven by a prevalence of insertions over deletions, reflecting population-level TE activity. Strikingly, both SVs and *de novo* TE insertions — driven by the concurrent mobilization of diverse TE families — segregate among *C. ariakensis* populations and contribute to genomic differentiation associated with local adaptation to contrasting sea salinity and temperature levels. Our study demonstrates the power of integrating long- and short-read sequencing to recover a high-confidence catalogue of SVs and *de novo* TE insertions, and provides empirical evidence that structural variation is an active evolutionary force generating potentially adaptive genetic variation in a key lineage of ecologically and economically important bivalves.

## Introduction

Structural variants (SVs) encompass a diverse range of genetic variants, including inversions, translocations, duplications, insertions, and deletions (longer than ≥ 50 bp), along with other complex genomic rearrangements, such as chromosomal fusions and fissions (Ho et al., 2020). While genetic and genomic studies have traditionally focused on Single Nucleotide Variants (SNVs), recent years have seen a growing body of evidence linking SVs to significant evolutionary processes (Hof et al., 2016; Berdan et al., 2021; Stuart et al., 2026). Indeed, SVs affect more base pairs than SNVs in the human genome (Frazer et al., 2009) and, more generally, they may represent the most important source of genetic variation between and within species (Wellenreuther et al., 2019). Therefore, integrating the analysis of SVs into population genomics studies, even in non-model species, is a fundamental step towards a deeper understanding of genome evolution.

While most SVs usually occur at low frequencies across populations, coherently with their generally neutral or deleterious effects (Zhou et al., 2019; Weissensteiner et al., 2020), multiple cases report their involvement in population differentiation (Kirkpatrick and Barton, 2006; Harewood et al., 2010; Hof et al., 2016). Transposable elements (TEs) are the richest source of SVs across many eukaryotes (Bourque et al., 2018). TE-related SVs not only include new TE insertions (i.e., TE presence/absence polymorphisms), but homologous TEs can also act as substrates for ectopic recombination and homology-dependent DNA repair mechanisms giving rise to further SVs, such as inversion, duplication, and deletion events (Balachandran et al., 2022). TE-derived SVs have been linked to the emergence of multiple novel phenotypes, such as the industrial melanism in the peppered moth (Hof et al., 2016), the loss of the tail in apes (Xia et al., 2024), and plumage patterns in birds (Weissensteiner et al., 2020, Lutgen et al., 2025), among others. Moreover, TE-derived variants can be especially important for rapid adaptations, due to the large amount of genetic variation that they can rapidly introduce (Stapley et al., 2015).

Oysters (order Ostreida) are a group of worldwide-distributed bivalves that include numerous important species for aquaculture. Numerous oysters, such as the Estuarine oyster *Crassostrea ariakensis*, also show high levels of physiological plasticity (Bromley et al., 2016) and experience wide ranges of different temperature and salinity, thus being able to adapt to highly dynamic environmental conditions (Zhou et al., 2003; Li et al., 2021a). Recent genome projects revealed that their genomes are highly heterozygous and with a high and diverse repetitive content (Peñaloza et al., 2021; Gundappa et al., 2022; Martelossi et al., 2023; Martelossi et al., 2024) and that structural variants might be important contributors to species differentiation (Qi et al., 2023). Moreover, high levels of within-individual structural variation between homologues chromosomes have also been observed both in oysters as well as in other bivalves with most of these variants related to TEs and leading to high levels of hemizygosity (*i.e.*, a haploid genomic region in a diploid individual; Gerdol et al., 2020; Calcino et al., 2021; Takeuchi et al., 2022; Liu et al., 2025). However, the potential role of TE insertions or TE-related genomic deletions in driving this haplotypic diversity, and the resulting impact of such variation on population differentiation and local adaptation, remains uninvestigated.

Here, we used long read genomes of four oyster species representative to two distinct genera (*C. ariakensis*, *Crassostrea gigas*, *Ostrea denselamellosa*, *Ostrea edulis*) to understand (i) the levels of structural heterozygosity in oyster genomes (ii) their relationship with TE activity and (iii) the impact of SVs and TEs in population differentiation and local adaptation to different sea salinity and temperatures in the estuarine oyster (**Fig. 1**). For these purposes, we took into careful consideration the strengths and limitations of SV calling softwares both when using long- and short-read datasets, benchmarking our results with simulations and selecting the most appropriate method. Finally, we showcase how heterozygous TE insertions can be confidentially identified in long-read-based haploid representations of diploid genomes, predicting active TEs at the population level and therefore facilitating hypothesis-driven research.

**Figure 1.**
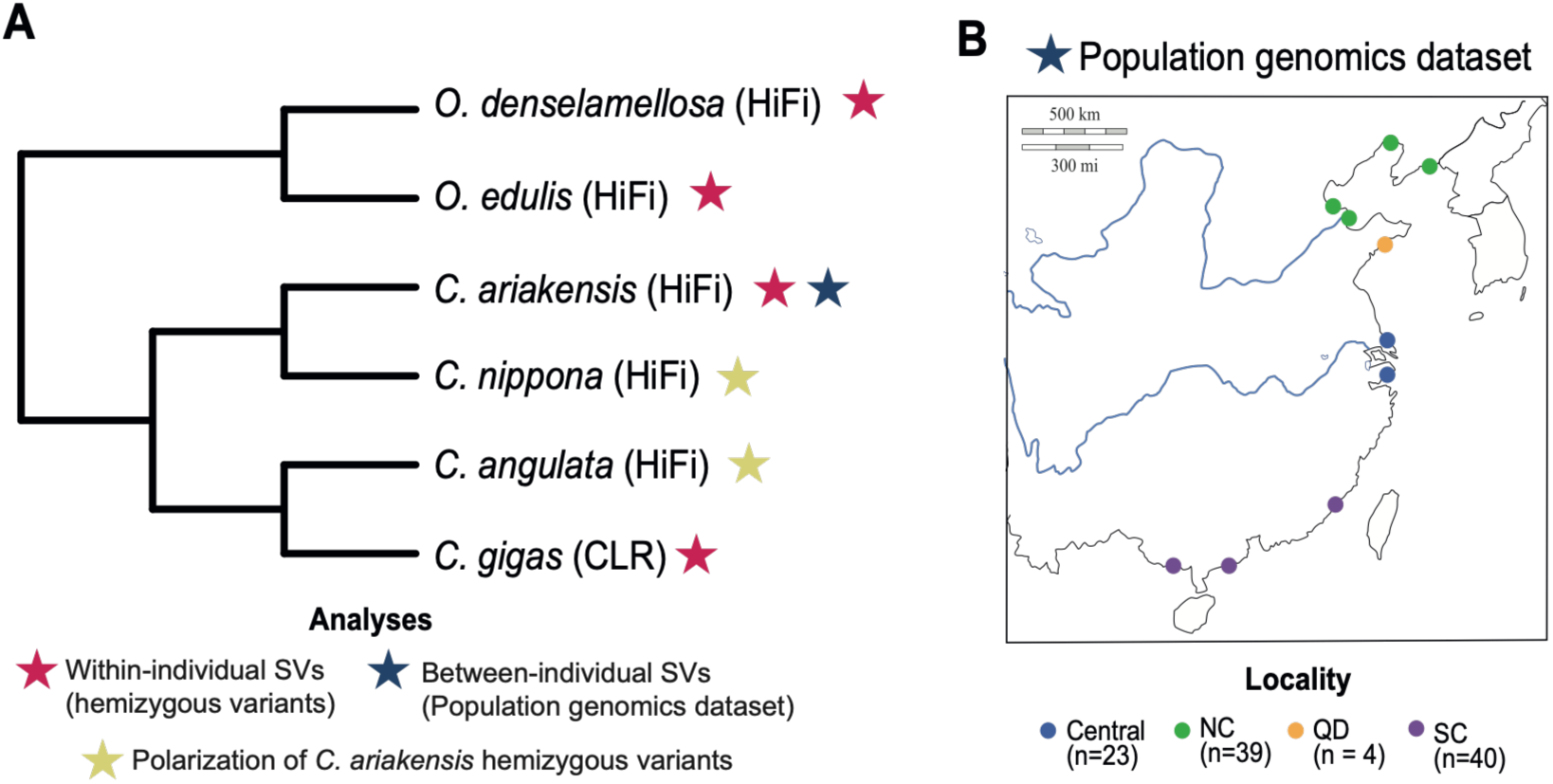
| Representation of publicly available genomic data used in this study. **(A)** Phylogenetic relationships of species included in all genomic analyses. For each species, the sequencing technology is represented, with stars indicating the type of analyses in which it was included. For within-individual SVs, see Material and Methods: “Heterozygosity prediction”; for between-individual SVs, see Material and Methods: “Between-individual SNV and SV calling”; and for polarization of *C. ariakensis* hemizygous variants, see Material and Methods: “Polarization of hemizygous variants and relationship with population-level TE activity”. **(B)** Geographic distribution of 107 estuarine oyster (*C. ariakensis*) WGA Illumina short-read re-sequencing samples included in population genomic and between-individual SV analyses. Sequencing data are from Li et al. (2021b).

## Materials and Methods

### Genomic datasets

We selected four publicly available high-quality oyster genomes to study their level of haplotypic variability: two *Crassostrea* (*C. gigas*, *C. ariakensis*) and two *Ostrea* (*O. edulis* and *O. denselamellosa*) species where a primary assembly is available online (**Sup. Tab. 1**; **Fig. 1A**). We chose only assemblies for which a wild or farmed but not inbred sample was sequenced using both long- and short-read technologies. Three of these genomes were sequenced with PacBio Hi-Fi technology: *C. ariakensis (*Wu et al., 2022), *O. denselamellosa* (Dong et al., 2023)*, O. edulis* (Adkins et al., 2023), whereas the *C. gigas* genome was sequenced with PacBio CLR (Peñaloza et al., 2021) reads. Additionally, we used PacBio HiFi reads from other two *Crassostrea* species, *C. nippona* (SRR23945211) and *C. angulata* (Teng et al., 2023; SRR20742558), to polarize the structural variants identified in the *C. ariakensis* genome (See Material and Methods: Estimation of genomic content and origin of hemizygous genomic regions; **Fig. 1A**). For population genomics and polymorphic TE insertion analyses, we used a recent dataset comprising 106 *Crassostrea ariakensis* genomes (NCBI bioproject: PRJNA715058) re-sequenced with paired-end Illumina short reads with a read length of 150 bp at a mean coverage of ∼20X (**Sup. Tab. 2**) and covering most of its distribution range (Li et al., 2021b; **Fig. 1B**).

### Transposable element annotation

For each genome, we generated a *de novo*, species-specific repeat library with RepeatModeler2 and the LTR pipeline extension (Flynn et al., 2020). To remove potential host genes, we built an oyster reference proteome database devoid of TE-derived proteins by combining the RefSeq annotations of *C. gigas* and *O. edulis* and filtering sequences showing BLASTP homology (E-value ≤ 1E−05; Altschul et al., 1990) to the RepeatPep database (Tarailo-Graovac & Chen, 2009). Each species-specific repeat library was then queried against this filtered proteome using BLASTX, and the results were processed with ProtExcluder. Cleaned libraries were merged with the Mollusca RepBase v20181026 and a set of manually curated bivalve consensus sequences (Martelossi et al., 2023; 2024). Redundancy was removed using CD-HIT-EST (Fu et al., 2012) with the 80–80–80 rule (≥80 % identity over ≥80 % of the shortest sequence for repeats ≥80 bp; Wicker et al., 2007). Finally, RepeatMasker (Tarailo-Graovac & Chen, 2009) was run in sensitive mode to annotate repetitive elements in the four genomes with the merged, non-redundant library.

### Heterozygosity prediction

For each genome, we calculated three distinct metrics of heterozygosity: (1) kmer-based heterozygosity, (2) SNP-based heterozygosity, and (3) structural heterozygosity, focusing on insertions and deletions (INDELs) between homologous chromosomes and therefore corresponding to hemizygous genomic regions.

For both kmer-based and SNP-based heterozygosity, we first cleaned Illumina short reads using bbduk (Bushnell, 2014), with a minimum quality threshold of 30 and a minimum length requirement of 35 bp. The filtered reads were then mapped to the genome using bwa-mem (Li, 2013), and their genomic coverage was determined using Mosdepth (Pedersen and Quinlan, 2018). Subsequently, we extracted the mapped reads and generated a kmer histogram with Jellyfish (Marçais and Kingsford, 2011), which was uploaded to Genomescope2 (Ranallo-Benavidez et al., 2020) for kmer-based heterozygosity estimation. SNPs were identified based on bwa alignments and bcftools mpileup (Li, 2011), retaining only biallelic variants with a genotype quality greater than 20, and called in genomic regions with coverage no greater than three times the median genome-wide estimation.

Structural heterozygosity, *i.e.* heterozygous INDELs along the whole genomes, was calculated based on the alignment of the long reads used for genome assembly against the assembly itself. For each genome, two alignments were generated: (1) using the minimap2 (Li, 2018) wrapper pbmm2 (https://github.com/PacificBiosciences/pbmm2) and (2) using LGNRM (Sedlazeck et al., 2018). In all instances, the appropriate preset option was selected based on the type of reads. SVs were then called for each alignment using the PacBio variant caller pbsv (https://github.com/PacificBiosciences/pbsv) and Sniffles2 (Smolka et al., 2024), providing genomic regions corresponding to tandem repeats as predicted by RepeatMasker. This resulted in four sets of SV calls, which were then filtered and merged to obtain a filter consensus set of reliable SVs. Firstly, variants not labelled as PASS and genotyped as homozygous for the alternative allele were removed, as they likely represented assembly errors or false positives. SURVIVOR (Jeffares et al., 2017) was used to: (1) retain variants supported by at least three SV sets (with a maximal distance of 1 kb between breakpoints, considering the SV type and its strand), (2) with a length greater than 49 bp, (3) supported by at least four reads, (4) at least 1 kb away from assembly gaps or ends of scaffolds, and (5) corresponding to INDEL events only. From the resulting VCF, an additional variant set was generated, preserving the same SVs but with breakpoints forced as estimated by pbsv instead of those chosen by SURVIVOR. Based on our benchmark results for all downstream analyses, we used the SV consensus set with breakpoints estimated by pbsv.

### SV calling benchmark

We ran a comprehensive benchmark of our SV calling pipeline using two distinct approaches: simulations with Sim-it (Dierckxsens et al., 2021) and short-read-based genotyping of previously identified hemizygous genomic regions with Paragraph (Chen et al., 2019).

For the simulations, we initially generated a synthetic haploid assembly of *C. ariakensis*, introducing 1,000 random deletions selected from the merged set with pbsv breakpoints inferred from the original genome. From this simulated assembly, synthetic Hi-Fi and CLR reads were generated at 15X genome-wide coverage, each with the appropriate technology-specific error profile. Simultaneously, synthetic long reads at 15X coverage were generated from the original genome, and the two read sets were merged. This approach allowed us to produce a true set of hemizygous deletions relative to the original genome and the reads carrying them. The merged reads were then mapped to the original *C. ariakensis* genome, and SVs were called using the same pipeline previously described. Bedtools *intersect* (Quinlan and Hall, 2010) was used to assess the reciprocal overlap (RO) between the true set and the called set. We applied three RO thresholds to consider a variant as correctly called: 80%, 90%, and 99%. A graphical representation of the simulation process is available in **Sup. Fig. 1**.

Additionally, we re-genotyped all INDELs of all genomes using the previously mapped Illumina short reads and Paragraph. Specifically, we re-genotyped the two consensus SV sets (i.e., with default SURVIVOR breakpoints and forcing pbsv-inferred ones) and considered a SV correctly called when also Paragraph genotyped it as heterozygous.

### Estimation of genomic content and origin of hemizygous genomic regions

We assessed the overlap between hemizygous deletions with transposable elements (TEs) as annotated by RepeatMasker in all four genomes. We considered a variant as TE-derived when it overlapped with an annotated TE for at least 70% of its length. To statistically test the overrepresentation of TE-derived variants in hemizygous deletions, we compared their observed number to a null distribution generated by randomly reshuffling all hemizygous deletions 10,000 times (excluding assembly gaps) and counting at each iteration the number of TE-derived variants. Furthermore, because deletions in the reference genome — in our case corresponding to the assembled haplotype — can represent both deletions in the other haplotype or insertions in the assembled one, we polarized the variants of *C. ariakensis* using PacBio HiFi reads from the two previously described *Crassostrea* species as reference (*C. angulata* and *C. nippona*). Briefly, hemizygous deletions of *C. ariakensis* were genotyped with SVJedy-graph (Romain and Lemaitre, 2023) and SNIFFLES2, and only variants with concordant genotype across at least three combinations of species and genotype were considered. When a *C. ariakensis* hemizygous deletion was genotyped as homozygous for the reference allele in *C. gigas*, it was considered a derived deletion event and an insertion event when it was genotyped as homozygous for the alternative allele.

### Polymorphic transposable element insertion analyses

To identify mobile element insertions across the 106 *C. ariakensis* samples we used TEMP2 (Yu et al., 2021). Briefly, Illumina short reads were mapped to the reference genome with bwa-mem and TEMP2 was run with default parameters. Results were filtered keeping only insertions supported by at least 10% of the median genome-wide read coverage, with read support at both ends and with a frequency in the sequenced genome ≥ 0.2. TE insertions were clustered between samples using bedtools *cluster* with a maximum allowed distance between their ends of 50 bp. TE insertions are considered to be virtually homoplasy-free variants (Okada et al., 2004) and we therefore considered two or more clustered insertions to derive from the same transposition event when involving the same TE consensus sequence (unique TE insertions). We classified unique TE insertions based on their frequency across samples of each *C. ariakensis* population: low-frequency when they appeared in 1 to 10 samples, intermediate-frequency appeared in 10 to 20 samples, and high-frequency s appeared in more than 20 samples, which corresponds to more than 50% of the individuals for both populations.

### Between-individual SNV and SV callings

Single nucleotide variants (SNVs) and small indels (length < 50 bp) were jointly called across the *C. ariakensis* population dataset using Platypus (Rimmer et al., 2014), retaining only variants with a minimum mapping and base quality of 20. From the resulting VCF file, we only kept biallelic SNVs called on the 10 *C. ariakensis* assembled pseudo-chromosomes and marked as PASS. Between-individual SVs were initially called independently in each sample using Manta (Chen et al., 2016) with default parameters. Sample-specific VCF files were then filtered, retaining only variants marked as PASS, and subsequently merged into a multi-sample VCF using Jasmine (Kirsche et al., 2023), allowing a maximal Euclidean distance of 20 bp between breakpoint representations of the variants. Paragraph was then used to re-genotype the merged VCF file for each sample to obtain population-scale genotyping, skipping genomic regions with a coverage higher than 20 times the median sample coverage, as recommended by Paragraph developers. The resulting multi-sample VCF file was further filtered, retaining only INDELs. Genotypes not marked as PASS by Paragraph were set as missing, and variants falling within tandem repeats were removed. Furthermore, we also removed variants genotyped as homozygous for the reference allele across all samples. Indeed, due to the strict cutoff in terms of distance between breakpoints applied during the SV merging process by Jasmine, redundancy in variants is expected in our merged VCF. Nevertheless, owing to the high sensitivity of Paragraph in breakpoint deviations (refer to Chen et al., 2019), redundantly incorrect representations are expected to be genotyped as 0/0 in all samples, thereby ensuring the accurate genotyping only of their best representation. Finally, similarly to what applied on SNVs, we only kept variants called on assembled pseudo-chromosomes.

### Population genomic analyses

We ran population genomic analyses similarly for both SNVs and SVs datasets, using only variants with a minor allele frequency (MAF) ≥ 0.05 and genotyped in at least 30% of the samples. PCA analyses were carried using PLINK (Chang et al., 2015) with default parameters. For SVs analyses, we analyzed both insertions and deletions separately and together. Population structure was inferred using fastStructure (Raj et al., 2014) with a parameter K (number of populations) ranging from 1 to 5. Due to the overlapping population structure recovered with PCA analyses on the three SVs sets, we only analyzed insertions and deletions together. The optimal number of populations was chosen based on their marginal likelihood, as calculated by the chooseK function from the fastStructure package. To identify candidate regions under selection between the north and south populations, we calculated fixation (*F_ST_*) statistics in a 100-kb sliding window with a step size of 10 kb, similarly to Li et al. (2021b). Genomic regions showing strong selection signals were defined as regions with the top 2.5% *F_ST_*. To identify genes recovered under selection between north and south *C. ariakensis* populations by Li et al. (2021b), we used Liftoff (Shumate and Salzberg, 2021) to lift their gene annotation obtained on a different *C. ariakensis* individual sequenced with Nanopore to the PacBio HiFi assembly. The lifted annotation was further used for functional annotation with eggNOG-mapper (Cantalapiedra et al., 2021) and GO terms enrichment analyses for genes within genomic regions under putative selection was performed with the topGO R package (Alexa and Rahnenführer 2025).

## Results

### Simulation-based benchmark of long-read based identification of hemizygous genomic regions

We combined the use of two different aligners with two distinct SV callers to obtain a reliable consensus set of heterozygous INDELs on a reference genome representing hemizygous genomic regions. Four genome assemblies from four oyster species, representative of two genera and two different PacBio sequencing technologies, were assayed: *C. gigas* (CLR), *C. ariakensis* (HiFi), *O. denselamellosa* (HiFi) and *O. edulis* (HiFi). Each combination of SV caller and long-read mapper produced different SV sets, with a number of variants ranging from 2,799 and 24,521 (**Sup. Fig. 2**). We therefore benchmarked the performance of each combination of methods using a simulated *C. ariakensis* genome where we introduced a true set of hemizygous genomic regions. The two-parameters precision (the proportion of true calls among all detected variants) and recall (the proportion of correctly identified variants among all true variants) estimated for each method combination, highlighted the strong influence of sequencing technology and SV calling pipeline on the accurate detection of heterozygous INDELs, especially under stringent RO thresholds between true and called variants. (**Fig. 2A-B**). As expected, HiFi reads consistently outperformed CLR reads in terms of both precision and recall rates, regardless of the read mapper and SV caller used. Requiring support from at least three callers (*i.e.*, relying on a consensus set) increased the precision rate but decreased the recall rate when using CLR reads (**Fig. 2A-1B**), whereas this drop was not observed when using HiFi reads. Sniffles2 outperformed pbsv in terms of precision, independently from the underlying aligner; however, pbsv achieved the highest recalling rates, especially at the stringent 99% RO requirement. These results may reflect a more accurate representation of SVs and their breakpoints by pbsv.

**Figure 2.**
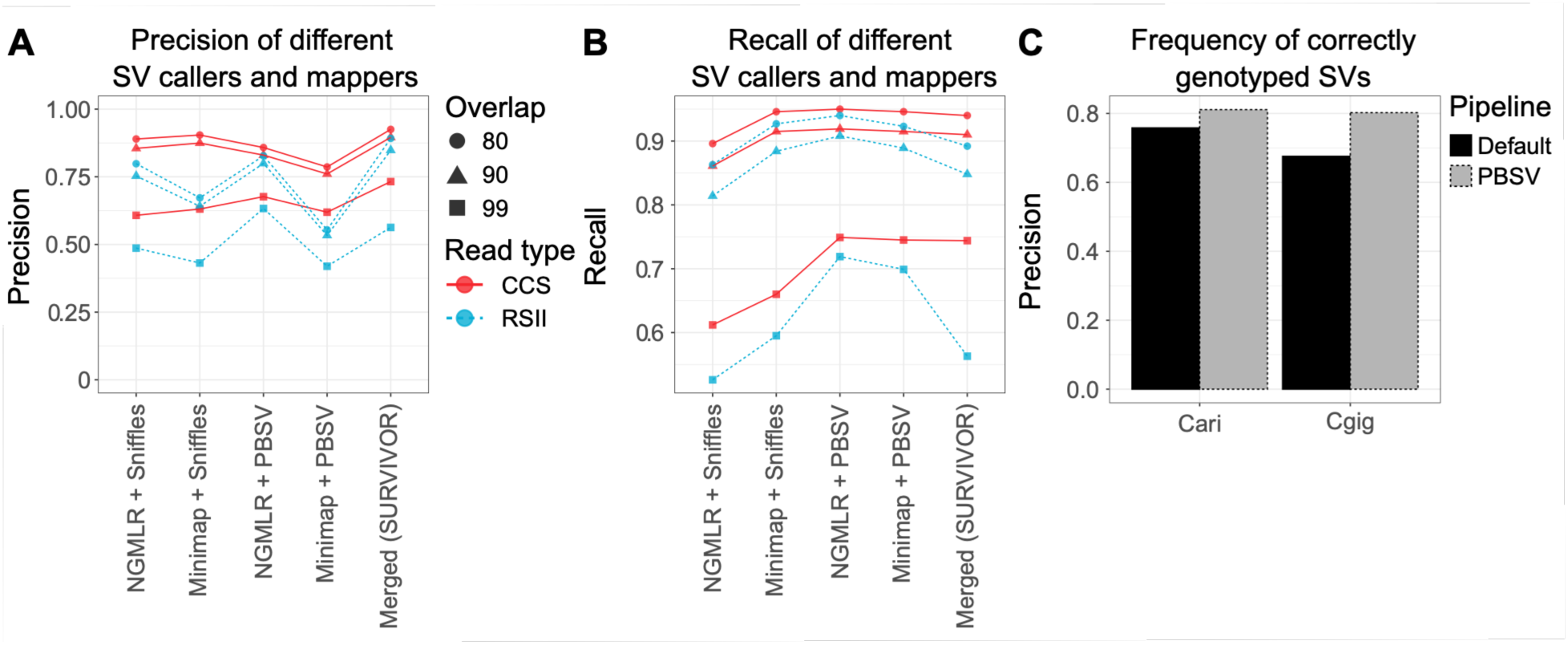
| Benchmark results of the SV calling pipelines using simulation and short read based re-genotyping of hemizygous variants. **(A)** Precision and **(B)** recalling rates based on simulations of 1,000 hemizygous variants in the *C. ariakensis* genome. **(C)** Frequency of correctly re-genotyped variants using short reads and Paragraph. Cgig = *C. gigas*, Cari = *C. ariakensis*.

To test this hypothesis, we considered the consensus set of heterozygous INDELs identified in *C. gigas* and *C. ariakensis* as a true set, and re-genotyped this set with Paragraph using Illumina PE reads sequenced from the same samples employed for long-read sequencing. Since Paragraph realigns reads to a sequence graph using stringent parameters during the genotyping process and is highly sensitive to breakpoint estimates (Chen et al., 2019), we expected a higher number of correctly re-genotyped variants when forcing pbsv SV breakpoints. Indeed, when these were compared to those automatically selected by SURVIVOR, we consistently observed an increase in the number of recalled variants, which always exceeded the 75% (**Fig. 2C**). In this case as well, we observed an impact of the sequencing technology: heterozygous INDELs identified in *C. ariakensis* were more frequently correctly genotyped using short reads, with a percentage of correctly re-genotyped variants equal to 81% and 84% for default and pbsv-forced breakpoint representation, respectively. *C. gigas*, for which CLR reads were used, benefited more from pbsv-estimated breakpoints, with an increase in the Paragraph re-genotyping rate from 68% to 80%. Thus, since a variant supported by at least three callers always included a pbsv-derived representation, we decided to rely on the SV consensus set with the breakpoints estimated by pbsv.

### Structural heterozygosity in oysters

To understand what types of genetic variants drive the heterozygosity in oyster genomes, we estimated and compared heterozygosity based on *k*-mer diversity, on the proportion of genomes involved in SNVs and percentage of genome in hemizygosity (structural heterozygosity). *K*-mer-based analyses estimated oyster heterozygosity to range from 0.8% for *O. denselamellosa* to 3% for *C. gigas* (**Fig. 3A**). SNV-based heterozygosity estimation consistently yielded lower values for all species, ranging from 0.6% for *O. denselamellosa* to 1.3% for *C. gigas*. Conversely, we observed high levels of structural heterozygosity, ranging from 3% for *O. edulis* to 13% for *C. ariakensis*. Given that we only considered INDEL between homologous chromosomes, this indicates that a significant proportion of the oyster genome exists in a hemizygous state (*i.e.*, a haploid genomic region in a diploid individual; **Fig. 3B**). *K*-mer and short-read counts across hemizygous genomic regions reflect what we may expect from genomic regions that are present in a single copy in a diploid genome: Indeed, contrary to the two-peak plots of the whole genome where both heterozygous and homozygous peaks are present, hemizygous regions only show a single peak that overlaps with the heterozygous peak of the whole-genome plot (**Fig. 3C**, **Sup. Fig. 3**).

**Figure 3:**
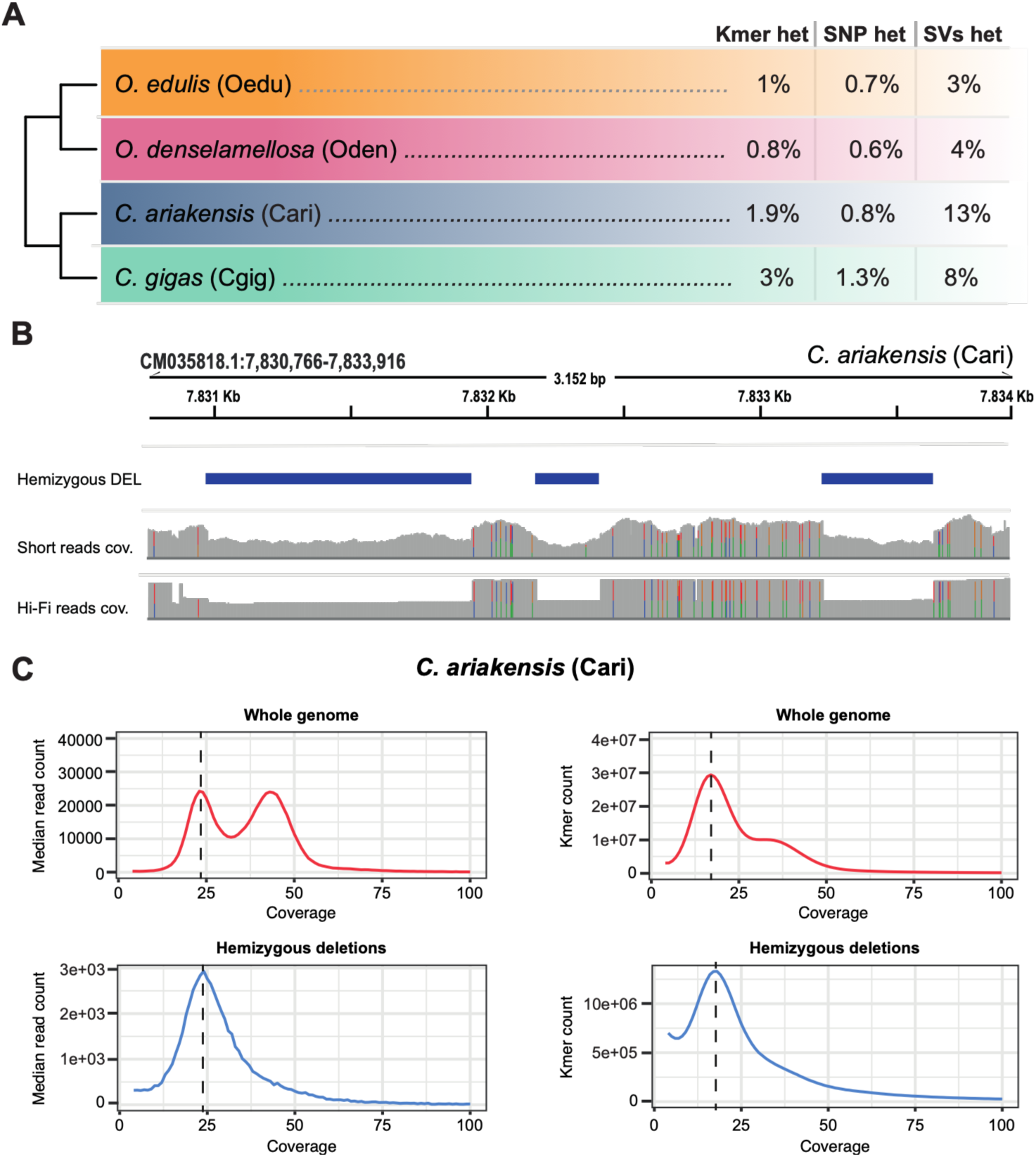
Heterozygosity estimations in four oyster genomes. (**A**) Heterozygosity levels estimated across the four analyzed oyster genomes. Kmer het = Kmer-based heterozygosity estimation, SNP het = SNP based heterozygosity, SV het = Structural heterozygosity in terms of insertions and deletions between homologous chromosomes and thus corresponding to hemizygous genomic regions. (**B**) Example of a *C. ariakensis* genomic region characterized by three hemizygous deletions (Hemizygous DEL). These regions show about half of the coverage of flanking sequences and are depleted of any SNP. (**C**) Median read count of 1-kb sliding windows and kmer count of all mapped reads in the *C. ariakensis* genome (red) and across hemizygous deletions (blue). The dotted line represents the heterozygous peak of the whole genome. Plots for all other species are reported in **Sup. Fig. 3**.

Short to medium-size INDEL (from 50 bp to 1 kb) contribute most to the total number of hemizygous variants, comprising 79%, 75%, 83%, and 85% of the total number of variants in *C. gigas*, *C. ariakensis*, *O. edulis*, and *O. denselamellosa*, respectively (**Fig. 4A).** TE-related variants, defined as those having an overlap of at least 70% with an annotated TE, represent 62%, 64%, 47%, and 52% of the heterozygous deletions in the same four species (**Fig. 4A**). In the *Crassostrea* samples, most of the TE-related variants are DNA TIR transposons (46% and 48% in *C. gigas* and *C. ariakensis*, respectively), followed by RC TEs (23% and 24%, respectively) (**Fig. 4A**). However, RC TEs outnumber DNA TIR transposons in the 2–4 kb size bin, where they account for 43% and 58% of TE-related variants in *C. gigas* and *C. ariakensis*, respectively, consistent with the high number of non-autonomous RC elements (mostly *Helentron-associated INterspersed Elements* - HINE) present in their genomes (Peñaloza et al., 2021). Within the *Ostrea* samples, where fewer TE-related variants were identified, we observed a less pronounced predominance of major TE groups (**Sup. Tab. 3**). For example, SINEs and LTRs in *O. denselamellosa* are predominant elements in the 200-400 bp size bin, constituting 32% and 24% of the variants, respectively. Due to the appreciable number of TE-related variants identified, we sought to determine whether hemizygous genomic regions are more likely to be associated with TEs compared to a null expectation. Permutation testing (10,000 reshuffles) revealed a significant overrepresentation of TE-related variants across all four species compared to the null distribution (**Fig. 4B**). This reinforces the role of TEs as a primary driver of within-individual structural variability.

**Figure 4.**
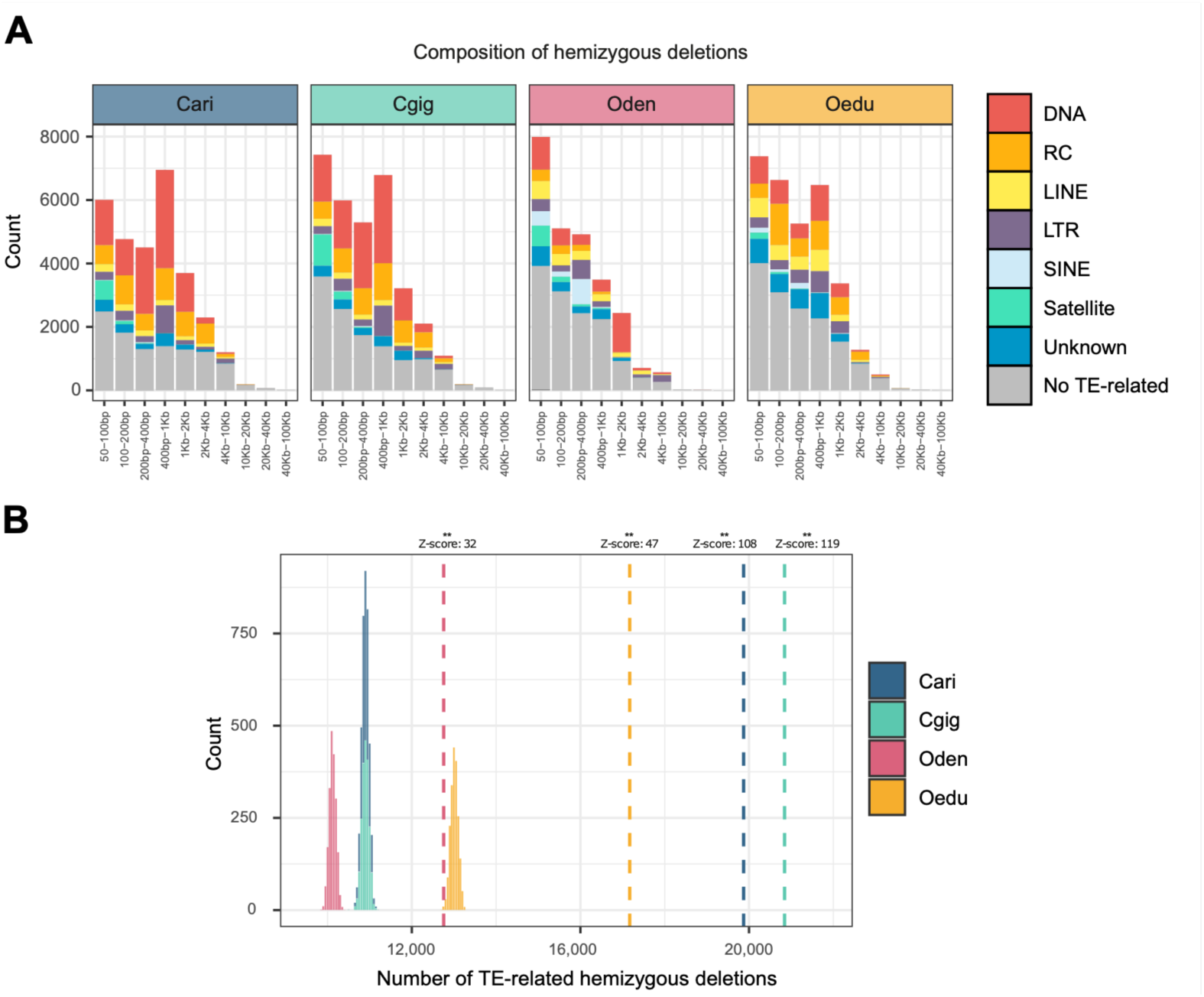
| Relationship between transposable elements and hemizygous variants. (**A**) Number of hemizygous deletions subdivided into different size bins and coloured by different TE types when classified as TE-related, defined as variants with an overlap with a TE annotation of at least 70%. (**B**) Null distributions of the number of TE-related hemizygous deletions generated by randomly reshuffling the variants 10,000 times across the genome and counting, at each iteration, the number of overlaps with a TE annotation, requiring at least 70% overlap. Dotted lines represent the observed number of TE-related variants. The Z-score of the observed number of TE-related variants compared to the null distribution is reported for each species. ** = p-value < 0.001. Cari = *C. ariakensis*, Cgig = *C. gigas*, Oden = *O. denselamellosa*, Oedu = *O. edulis*.

### Polarization of hemizygous variants and relationship with population-level TE activity

Hemizygous genomic regions can arise due to a novel insertion or the deletion of a genomic region. To assess the impact of insertion and deletion events on the emergence of hemizygous genomic regions in *C. ariakensis*, we polarized all its deletions by genotyping with PacBio Hi-Fi reads from the two closely related species, using two different SVs genotypers: SVJedy-Graph and Sniffles2. Out of the 5,484 variants with a concordant genotype in at least three of the four different analyses, 965 (18%) were identified as homozygous for the reference allele, indicating potential deletion events in *C. ariakensis* (**Fig. 5A**). Another 56 (1.02%) were genotyped as heterozygous, suggesting shared hemizygous regions across all species or genotyping errors. The remaining 4,463 (81%) were genotyped as homozygous for the alternative allele, implying they are the result of *bona-fide* TE insertions present in an heterozygous state in the *C. ariakensis* reference genome. Coherently, most of these polarized insertions were found to be TE-related (72%). Transposons included in such variants appeared significantly younger based on their divergence from to the consensus (K2P distance) than those involved in deletion events (Wilcoxon rank-sum test with Bonferroni correction, p < 0.001; **Fig. 5B**). To assess whether heterozygous TE insertions on a reference genome may serve as indicators of TE activity across populations, we compared the transposon families contributing to the emergence of heterozygous TE insertions in the *C. ariakensis* genome with *de novo* TE insertions identified in 107 short read-sequenced, wild-caught individuals collected along the entire Chinese coastline (**Sup. Tab. PopGen Dataset**). We identified a total of 55,383 mobile element insertions, ranging from a minimum of 292 to a maximum of 706 per sample (mean = 530). These were clustered in 17,215 unique insertions between samples (**Fig. 5C**). Most of these are related to DNA TIR transposons (62%) with Crypton-A, Kolobok, PIF-Spy and TcMar-Pogo contributing between 1,127 and 443 insertion events (**Fig. 5C,Sup. Tab. 4**). Both RC/Helitron and LTRs constitute 17% of the total number of unique insertions (respectively 2,897 and 2,852s), with Gypsy elements that constitute the great majority of LTRs (1,982). Regarding LINEs (3.8% of the total number of unique insertions), we found that most derive from CR1-Zenon and RTE-X elements. Finally, very few SINE-related insertions were found (0.3%), with most of them coming from the tRNA-V (49) superfamily. Lastly, we observed a significant positive correlation between the number of unique insertions annotated for each family (i.e., consensus sequence) and the number of hemizygous genomic regions classified as insertions involving the same repeat family after polarizing the variants (Spearman’s rank correlation rho = 0.56, p-value < 0.01; **Fig. 5D**).

**Figure 5.**
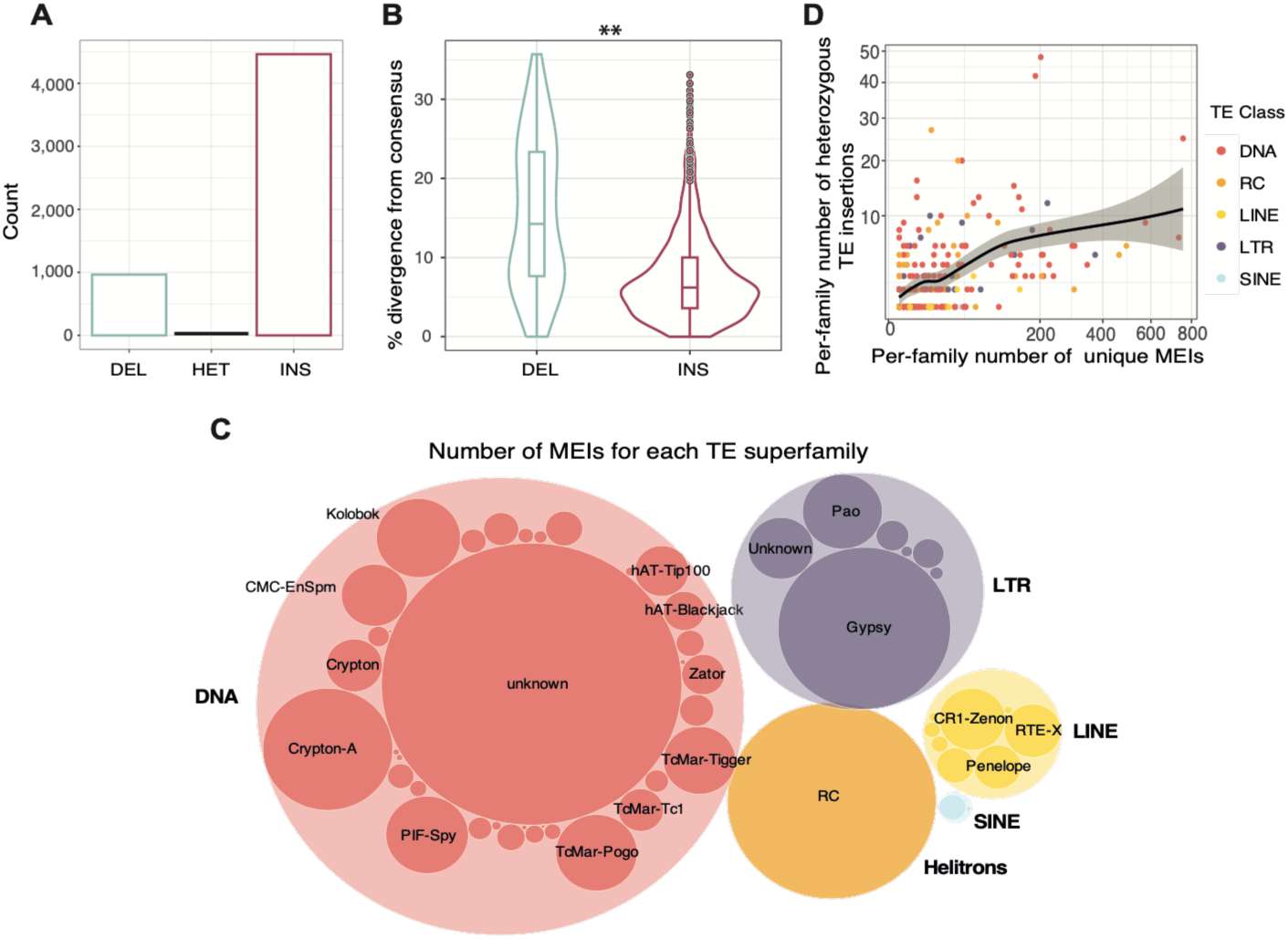
| Polarization of hemizygous variants and their relationship with polymorphic TE insertions the estuarine oyster. (**A**) Number of SVs identified in the reference genome of *C. ariakensis*, categorized as deletions, shared hemizygous genomic regions and insertions, as inferred based on their genotype in the closely related species *C. angulata* and *C. nippona*. (**B**) Divergence of TE copies compared to their consensus sequences expressed as K2P distance, used as a proxy for the relative age of the insertion. The two sets of values are significantly different (Wilcoxon rank-sum test with Bonferroni correction, p-value < 0.01). (**C**) Proportion of unique polymorphic TE insertions belonging to different repeat superfamilies identified in the estuarine oyster population genomic dataset. (**D**) Scatterplot depicting the per-family number of polymorphic TE insertions identified in the estuarine oyster population genomic dataset and the per-family number of heterozygous TE insertions identified in the *C. ariakensis* reference genome (Spearman’s rank correlation rho = 0.51, p-value < 0.01).

### Population genomic analyses based on SVs and mobile element insertions

Using the previously described population dataset of estuarine oyster (**Fig. 1B**), we identified a range of 3,092 to 15,024 insertions (mean = 7,940; sd = 3,348) and 18,151 to 32,876 deletions (mean = 25,434; sd = 3,521) per sample. After consolidating individual SV sets by performing joint genotyping with Paragraph, and filtering the resulting variants, we retained 255,140 SVs. The majority of these SVs correspond to deletions relative to the reference genome (83%) and 53% of these overlap with TEs for at least 70% of their length.

After retaining only SVs with a MAF of at least 0.05 and successfully genotyped in at least 30% of the samples, we used 74,216 deletions (69% of which overlapping with TEs), 29,687 insertions, and the joint dataset of insertions and deletions to infer population structure. PCA analyses based on all three variant sets mirror the original results based on SNVs obtained by Li et al., (2021b), as well as our re-analyses based on 1,285,295 SNVs (**Fig. 6A; Sup. Fig 5**), indicating the high quality of our population-informative SV set. In particular, the two main populations encompassing all northern (NC) and southern (SC) samples are clearly separated, with the presence of two additional clusters including all samples from central China, on one side, and individuals from the Qingdao locality (QD), on the other side. Admixture analyses based on the joint dataset indicated that the best-fitting number of population clusters was two with ancestry that reflects the PCA-based population structure (**Fig. 6B**).

**Figure 6.**
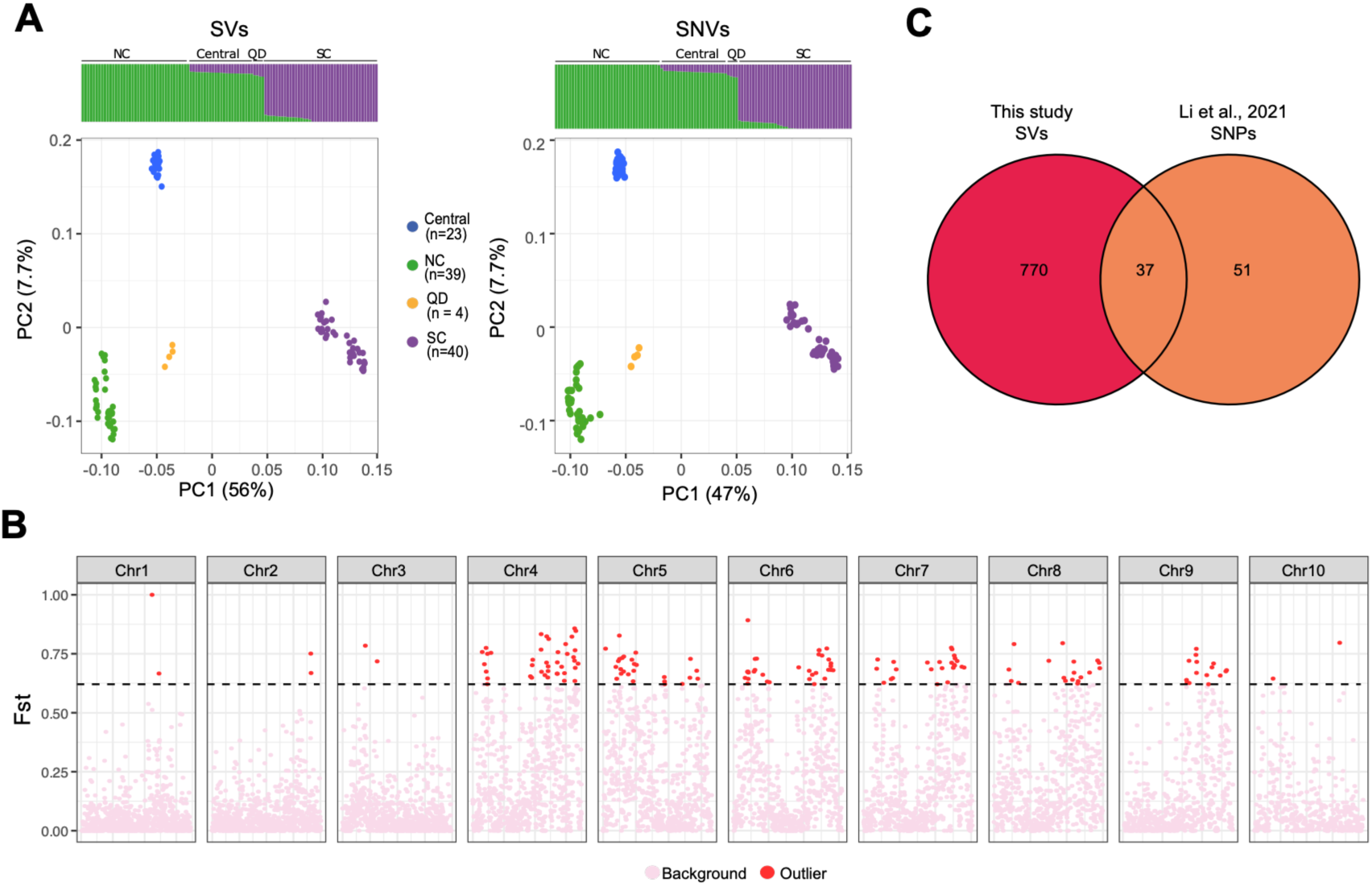
| Population genomics analyses of the estuarine oyster. (**A**) FastStructure and PCA analyses based on population informative SNVs and SVs. For the geographic distribution of the samples see Fig. 1B (**B**) Manhattan plot with SVs *F_ST_* values over 100-kb genomic sliding windows along the 10 *C. ariakensis* chromosomes. The dotted line shows the 97.5% interval of genome-wide values. Red circles are candidate genomic regions under divergent selection. (**C**) Overlap between genes under putative divergent selection in this study based on SV analyses and in Li et al. (2021b) based on SNVs.

Furthermore, we used *F_ST_* on the joint SV set to find regions putatively under selection, and identified 148 candidate regions between NC and SC populations (*F_ST_* > 0.62) encompassing all 10 chromosomes, for a total of 14.8 Mb and 770 genes (**Fig. 6C**). Twenty-one GO terms were found to be enriched among genes putatively under selection (**Sup. Tab. S5**), with several directly associated with ion transport, among which potassium ion import, (GO:0034762, GO:0043269), amino acid and lipid metabolism (GO:0006629,GO:0170044), negative regulation of TOR signaling (GO:0032007) and amino acidic catabolic processes (GO:0170044, GO:1901606, GO:0006567). Furthermore, these candidate genes included 37 out of the 51 loci previously identified by Li et al. (2021b) as being under selection based on SNV data (**Fig. 5D**). Among them, 14 genes were also found to be differentially expressed in oysters exposed to contrasting salinity and temperature conditions (**Sup. Tab. S6**), among which members of the solute carriers families *Slc23a2* (Solute carrier family 23 member 2) and *Mct12* (Monocarboxylate transporter 12).

When specifically examining mobile element insertions, we focused on NC and SC populations due to the high and similar number of samples, finding more insertions segregating in the NC population compared to SC samples (**Fig. 7A**). Most of them are present as singletons or at low frequencies across SC and NC populations (**Fig. 7A**). Indeed, among the 66 and 81 high frequency insertions within their geographical area, only eight and two, respectively, were found with a frequency equal to or greater than 90%. Also, the 2,550 insertions shared by the two populations in at least one individual (**Fig. 7B**) exhibited low frequencies (**Sup. Fig. 6**). However, when performing cluster analyses on MEI frequencies, we observed again a clear separation between SC and NC samples, with Central and QD samples nested within the latter (**Fig. 7C**). We identified a set of 18 mobile element insertions exhibiting high frequency in one population (NC or SC) but low frequency in the other, alongside 12 additional high-frequency insertions that were completely absent from the contrasting population (**Sup. Fig. 6**). These insertions involve DNA TIR elements (Crypton-A, PIF-Spy, PIF-Harbinger hAT-Ac and Academ-2), RC Helitrons, and Gypsy LTR transposons. In addition, 325 unique insertions were localized within the highly diverging regions defined by the *F_ST_* analyses of the SVs dataset. While most of these originated from TE superfamilies with differential frequencies across SC and NC populations (**Sup. Tab. S7**), only three insertions showed a significantly diverging frequency between the two populations (two RC/Helitrons and one DNA/Crypton-A element).

**Figure 7.**
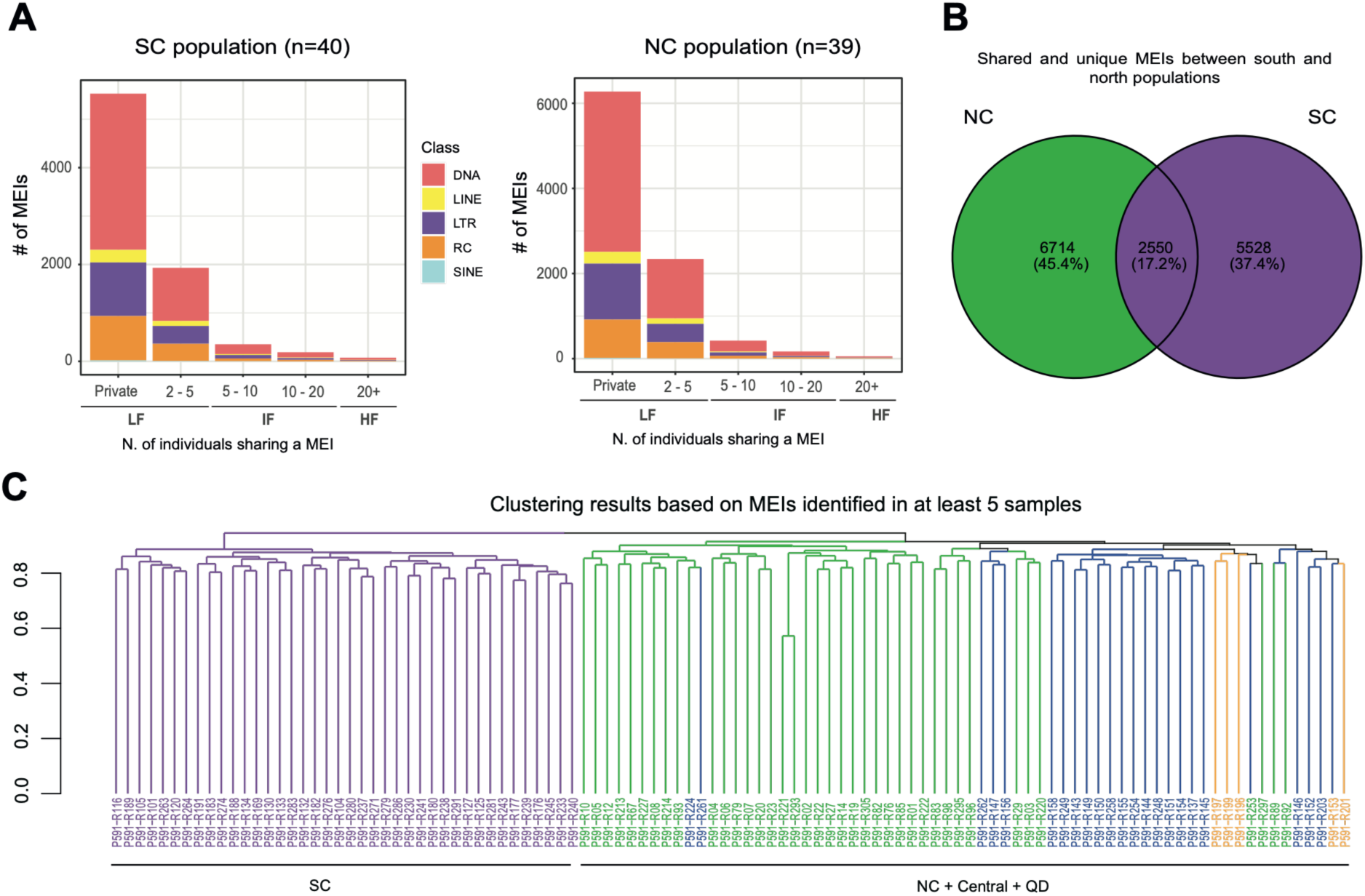
| Polymorphic TE insertion dynamics in the estuarine oyster. (**A**) Frequency of non-reference TE insertions across individuals of southern (SC) and northern (PC) populations. LF = Low frequency, IF = Intermediate frequency, HF = High frequency (**B**) Number of unique and shared insertions between *C. ariakensis* populations. NC = Northern China population; SC = Southern China population. (**C**) Clustering analyses based on non-reference TE insertions identified in at least five samples. Colors of tips and branches represent the geographic origin of the samples.

## Discussion

### Precise inference of SVs from long-read data and consensus call sets require careful choice of variant representation

Structural variant calling’s accuracy can greatly differ depending on the underlying sequencing technology, coverage and SV calling pipeline (Mahmoud et al., 2019; Dierckxsens et al., 2021). Therefore, to quantify our ability to detect heterozygous SVs on a reference genome sequenced with two different PacBio technologies, we benchmarked the performance of two different SV callers (pbsv and Sniffles2) and two long-read mappers (Minimap2 and NGLMR). Importantly, we here simulated two sequencing libraries at 30X coverage which represent the current standard for *de novo* genome assembly (Earth Biogenome Project assembly recommendations: https://www.earthbiogenome.org/report-on-assembly-recommendations#:~:text=We%20provide%20three%20recommended%20specific,ONT%2Dbased%2D2). We found that different combinations of aligners and SV callers can produce significantly different sets of SVs. Therefore, a consensus strategy is fundamental to reducing false positive rates (Mahmoud et al., 2019; Dierckxsens et al., 2021; Balachandran et al., 2022). On the other hand, consistent with previous benchmarks (Dierckxsens et al., 2021; Balachandran et al., 2022), we found that pbsv estimated more precise SV breakpoints than Sniffles2, regardless of the PacBio sequencing technology or long-read mapper used. More accurate breakpoint estimation can improve the genotyping of SVs (Romain and Lemaitre, 2023). In particular, studies that detect high-confidence SVs using long reads in a panel of species representative of one or more populations, and subsequently genotype these variants at large scale on short-read resequencing datasets to investigate their evolutionary dynamics and possible phenotypic effects (Stuart et al., 2026), can greatly benefit from such improvements. Moreover, precise breakpoint inference is essential when investigating the mechanisms underlying SV formation, as microhomologous regions shorter than 50 bp can be sufficient to trigger a wide range of complex SVs (Liu et al., 2012; Chen et al., 2023).

### High levels of structural heterozygosity in oysters derive from TE insertions and predict population-level TE activity

Applying our within-individual SV calling pipeline to four high-quality oyster genomes, we found high levels of structural heterozygosity: with between 3% to 4% of *Ostrea* spp. and 8% to 14% of *Crassostrea* spp. genomes being in a hemizygous state. Structural heterozygosity surpasses both *k*-mer-based and SNV-based heterozygosity estimates, indicating that long insertions and deletions contribute significantly more to haplotypic variability compared to SNVs. Similar results were obtained in previous estimations of hemizygosity in different bivalve species using a comparable approach. Calcino et al. (2021) estimated 10.68% of the *Pecten maximus* genome to be in a hemizygous state, while Takeuchi et al. (2022) estimated 18.12% of hemizygosity in the oyster *Pinctada fucata*. While both studies used a similar approach of re-mapping long-reads against the haploid representation of the primary assembly, they relied exclusively onpbsv. As previously discussed, without a consensus strategy, this may have led to inflated estimates due to the high false-positive rates inherent to single-tool analyses. In contrast, our estimations — which integrate multiple long-read aligners and SV callers — are more robust and likely provide a more accurate representation of INDEL events between haplotypes.On the other hand, a recent comparison of fully-phased assemblies across a wide range of different bivalve species revealed that 46%-84% of their genomes is highly divergent between the two haplotypes, contributing to gene expression and phenotypic plasticity (Liu et al., 2025). By comparison, highly diverging haplotypes account for only 2.16%, 2.33%, and 21.77% of the human, chicken, and zebrafish genomes, respectively (Liu et al., 2025). Our results therefore corroborate that bivalves possess an exceptionally dynamic genome, which has likely facilitated the origin and maintenance of high genetic diversity throughout their evolutionary history (Liu et al., 2025).

Across all the aforementioned studies, as well as in our results, TEs have been found to be significantly associated with hemizygous and highly diverging genomic regions between homologous chromosomes. However, the relative contribution of TE insertions versus deletions to this structural variability was previously undetermined. TEs are known to mediate genomic deletions via homologous and non-homologous recombination, which can result in the loss of both the TE and adjacent sequences (Bourque et al., 2018). Here, we polarized the variants identified in *C. ariakensis* with HiFi long reads from two other closely related oysters and found that most of them are *bona-fide* TE-derived insertions, with a little contribution of deletion events. This finding was further supported by the unimodal distribution of divergence from consensus sequences of these TEs centered at low divergence, indicating these are recent TE copies. On the other hand, TE-derived deletions show a much wider distribution of divergences. Indeed, deletion events are expected to randomly involve TE copies, either young or old (i.e., less or more divergent from the consensus). Though, this does not include DNA TIR transposons which are mobilized through cut-and-paste mechanisms: when mobilized, cut-and-paste TEs present in homozygosity are expected to generate both an heterozygous insertion at the recipient site and a heterozygous deletion at their donor genomic location (Chen et al., 1992). We consider it therefore likely that highly active DNA TIR transposons contribute to both our insertion and deletion estimates.

The identification of heterozygous SVs based on alignments of whole genomes or phased haplotypes has received much attention in the recent years to aid in the identification of recently active TEs (Groza et al., 2024; Quah et al., 2025; Qian et al., 2025) and in TE library construction (Baril and Croll 2023; Sierra and Durbin 2024; Wang et al., 2025). Here we show that the use of collapsed representations of diploid genomes and associated long-read data, can provide clues on which TE families are more likely to be active at the population level. Because the identification of polymorphic TE insertions within populations greatly benefits from high-quality TE consensus sequences (Suh et al., 2018; Carrasco-Valenzuela et al., 2025), this information can be used to prioritize manual curation efforts and facilitate hypothesis-driven research.

### Structural variants contribute to population differentiation and local adaptation in C. ariakensis

Given the high within-individual structural variability found in oyster genomes, we hypothesized that transposons, and structural variants in general, may contribute to genetic differentiation between oyster populations and particularly to the highly structured populations of the Chinese estuarine oyster (Li et al., 2021b). Here NC and SC populations are clearly separated in accordance with the summer ocean currents of the Chinese coast where northern and southern currents remain separated, and they only meet near the Yangtze River estuary, at the location of the Central population (Ni et al., 2017). Moreover, NC populations experience, on average, 10.98 ‰ higher salinity, whereas SC populations experience an average monthly sea surface temperature increase of 10.35 °C (Li et al., 2021b). Therefore, the estuarine oyster represents an ideal model to study the role of SVs in shaping adaptive evolution and particularly to the two main environmental factors sea temperatures and sea salinity. The results of admixture and PCA analyses based on the SVs dataset confirmed our hypothesis, since they almost perfectly match with one another and with previous results based either on few loci or whole genome SNV data (Xiao et al., 2010, Li et al., 2021b). The high concordance between the population structures recovered with both datasets also implies that, despite long reads being the gold standard for SV calling (Mahmoud et al., 2019), short reads can still be successfully used to obtain high-quality SV sets, at least for insertions and deletions (Weissensteiner et al. 2020), and should therefore be considered in population genomic studies of bivalves. We found that genomic regions with high *F_ST_* values between NC and SC populations calculated with the joint SVs set have a great overlap with the *F_ST_* peaks detected by Li et al. (2021b) based on SNV data, as shown by the common set of genes identified under putative selection. Many of these genes were previously found to be differentially expressed under contrasting thermal and salinity conditions, such as *Slc23a2* and *Mct12* (Li et al. 2021b; **Fig. 5D; Sup. Tab. S6**). Moreover, GO term enrichment analyses of genes overlapping with SVs revealed many enriched biological functions to the same two environmental factors. Ion channels are known to play a fundamental role during osmotic stress in bivalves by mediating the ionic steady state of Na+, Cl-, K+ and Ca2+ (Brown and Piscopo, 2011; Meng et al, 2013; Tomanek 2014; Zeng et al, 2024). Moreover, when the cell membrane is depolarized under stress conditions, voltage-gated Na+/K+ channels are co-regulated to maintain osmotic balance (Marom 1998). Thermal and salinity acclimatation also involved lipid metabolism affecting cellular membrane fluidity (Hazel, J. R. 1995; Pernet et al, 2007) and increased energy demands during the acclimatization process (Luvizotto-santos, 2003; Lin et al, 2025). Similarly, TOR complexes are serine/threonine kinases that regulate cell growth, metabolism, and energy homeostasis in response to nutrient availability, stress, and environmental cues (Wullschleger et al, 2006). Finally, intracellular free amino acids have been shown to play a significant part as intracellular osmoregulators (Baginski and Pierce, 1977; Kube et al., 2007; Meng et al. 2013). Amino acid catabolism may therefore play a key role in the turnover or remodeling of amino acids used as osmolytes.

One of the major impacts of SVs is their potential to reshape *cis-*regulatory elements nearby genes, as well as the overall 3D genome architecture modulating *trans* gene regulation (Chiang et al., 2017; Spielmann et al., 2018). In *C. ariakensis,* strong signatures of selections were identified along intergenic genomic regions between NC and SC populations. Moreover, gene expression plasticity seems to be involved in these adaptations (Li et al., 2021b). We therefore speculate that SVs are likely involved in these adaptive processes in *C. ariakensis* and potentially in other oyster species.

### TE activity in *C. ariakensis* populations

TEs are one of the major sources of SVs, both because of the insertions themselves and because their repetitiveness across the genome can promote deletions, translocations and inversions through non-allelic homologous (ectopic) recombination (Balachandran et al., 2022). In our SV datasets, we found that most insertions are related to TEs. This, together with the finding that most of the polarized hemizygous deletions in the reference *C. ariakensis* genome are due to *bona-fide* TE insertion events, led us to analyze them at the population level in a dedicated analysis of the estuarine oyster population genomic dataset. A high number of different TE superfamilies appear to contribute to mobile element insertions, confirming that oysters possess a highly diverse mobilome where multiple TE lineages are concurrently active. For example, we identified a high number of mobile element insertions for TcMar-Pogo, CMC-EnSpm and Kolobok DNA TIR transposons as well as for LINE RTE-X, L1-Tx1 and CR1-Zenon for which autonomous elements elements were identified in their genomes (Martelossi et al., 2023). Similarly, most of SINE insertions are related to the tRNA-V superfamily, coherently with its identification in the *C. ariakensis* genome (Martelossi et al, 2024).

While TE activity is generally characterized as neutral or deleterious, a growing body of empirical evidence suggests that individual TE insertions, much like other structural variants, can play a pivotal role in adaptive evolution (Pasyukova et al., 2004; Blumenstiel et al., 2014; Barrón et al., 2014; Rech et al., 2022). In line with the expectation of purifying selection against deleterious or nearly neutral insertions, we found that the majority of mobile element insertions are restricted to single individuals. Alternatively, this pattern may stem from novel insertions restricted to single individuals. Irrespective of the mechanisms underlying the low frequency of mobile element insertions, the distinct grouping of NC and SC populations observed in our cluster analysis indicates that the two populations possess divergent TE landscapes. Our investigation into mobile element insertions with contrasting frequencies between these groups revealed both population-restricted TE insertions and the differential retention of ancestral polymorphisms between the two populations. Consequently, both *de novo* transposition events and differential sorting of standing genetic variation might have been involved in adaptations to local environmental pressures.

## Conclusions

Here, we extensively characterized within- and between-individual SVs and their relationships with TEs in an economically important clade of bivalves. Firstly, we showed that TE insertions are primary contributors to haplotypic structural differences in oysters with little contribution of deletion events. Concurrently we demonstrate that heterozygous SVs can be confidently called on haploid representations of diploid genomes assembled from long-read technologies and can aid in the identification of potentially active TEs at the population level. Secondly, we showed that SVs are fundamental contributors to population differentiation and local adaptations to different sea salinity and temperature levels in the Chinese estuarine oyster, potentially shaping its regulatory landscape and promoting gene expression plasticity. Finally, we found that the highly dynamic genome of the eastern oyster is reshaped by the activity of a high number of different TE families, with TE insertions segregating within populations. Our findings corroborate oysters, and bivalves in general, as a promising model system to study TEs and SVs, providing an important example on how to implement SV and TE analyses in population genomic studies of non-model species.

## Author contributions

JM and FG conceived the study with inputs from VP, AS and AL. JM performed bioinformatic analyses with the supervision of VP and AS. JM and VP prepared the first draft and all figures and supplementary data. All authors critically revised the manuscript.

## Suplementary figures

**Sup. Fig. 1:**
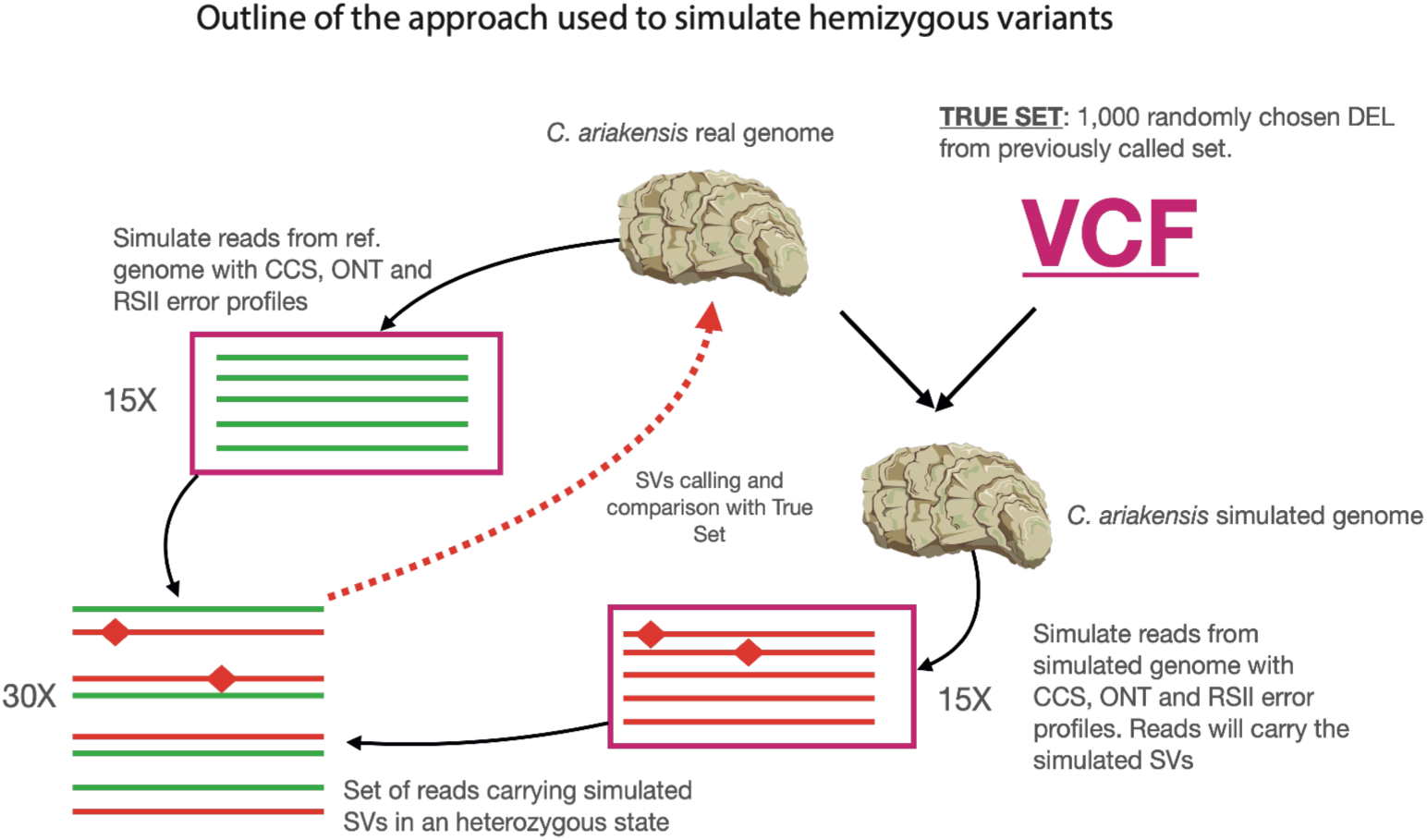
Schematic overview of the simulation approach used to evaluate the ability of different SV callers and long-read mappers to detect heterozygous SVs using different PacBio sequencing technologies.

**Sup. Fig. 2:**
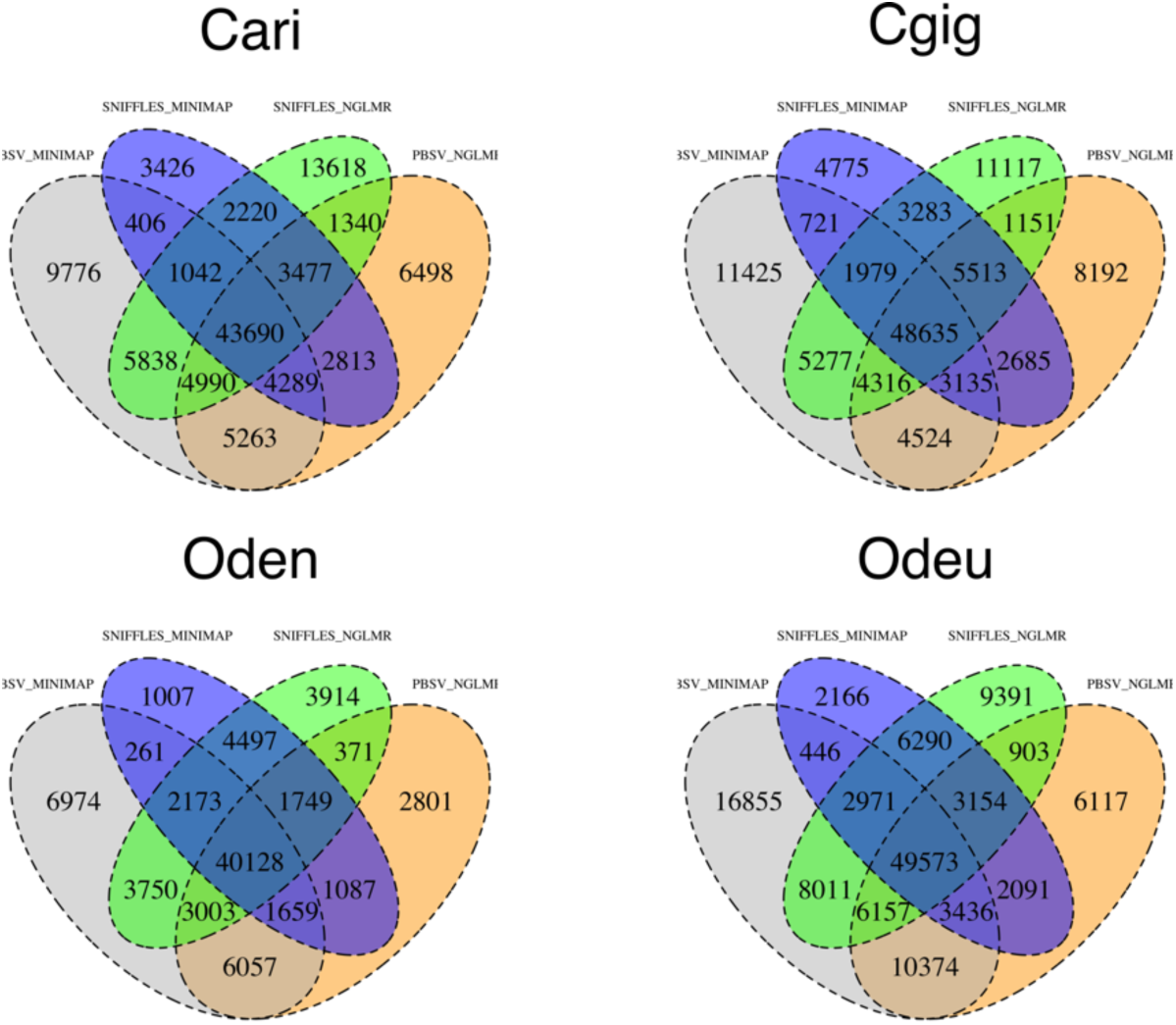
Number of unique and shared SVs called by different combinations of SV callers and long-read mappers.

**Sup. Fig. 3:**
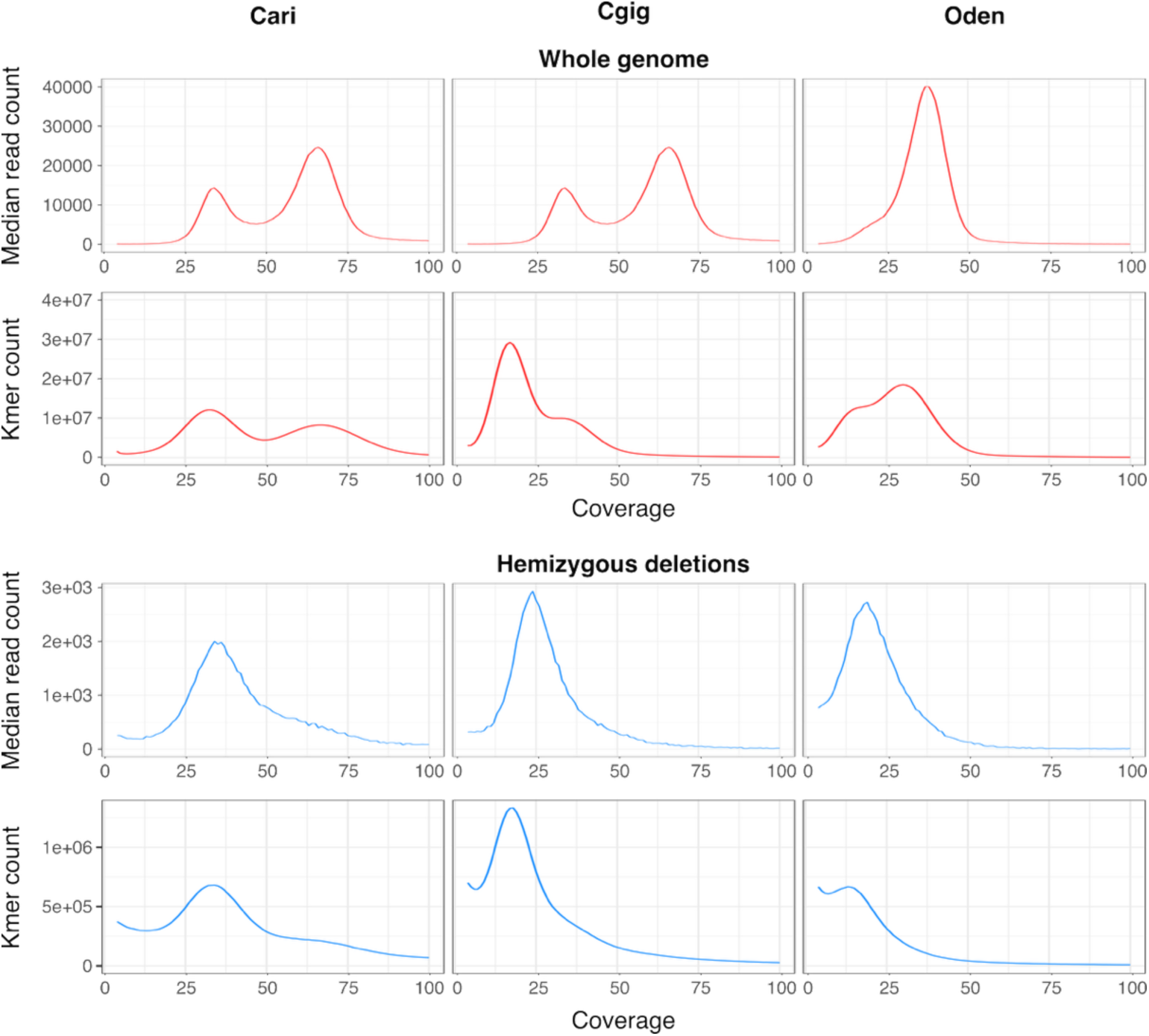
Kmer and short-read coverage along the whole genome and along only the identified homozygous deletions in the four analyzed oyster genomes.

**Sup. Fig. 4:**
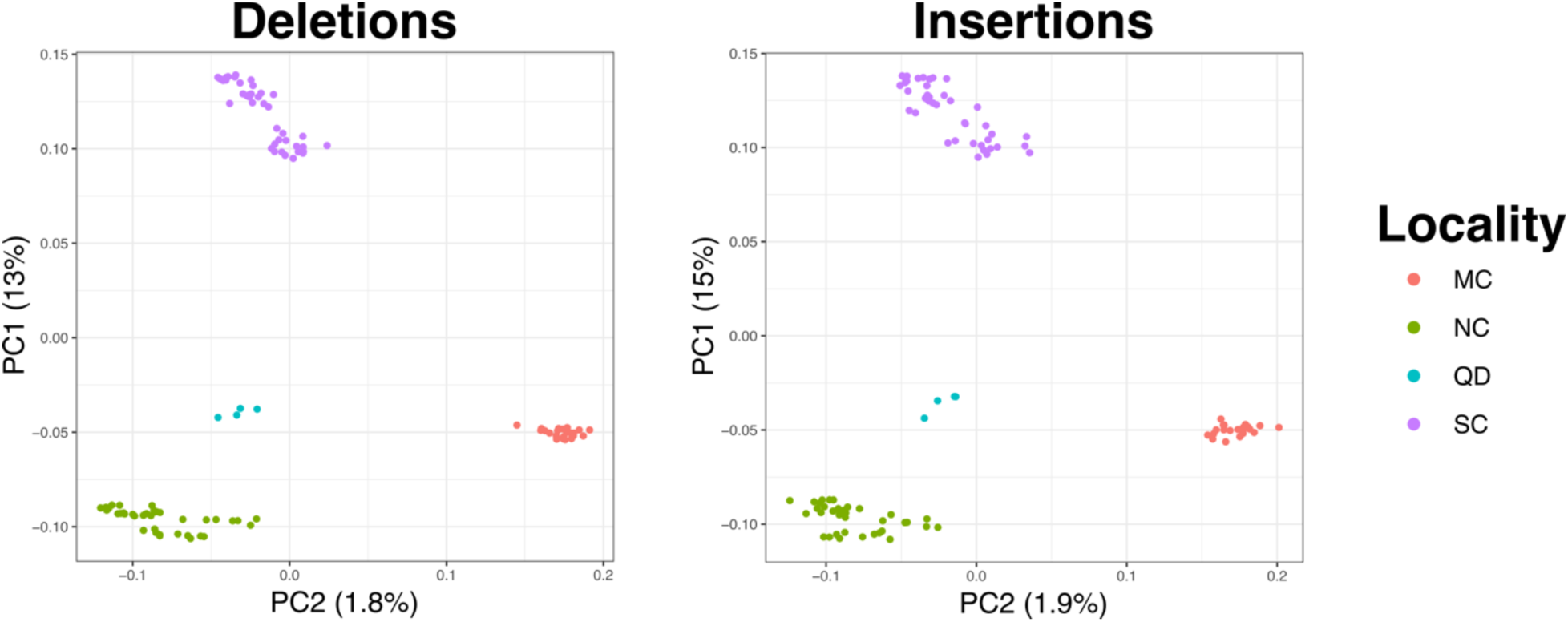
PCA analyses on the *C. ariakensis* population dataset based on insertions and deletions.

**Sup. Fig. 5:**
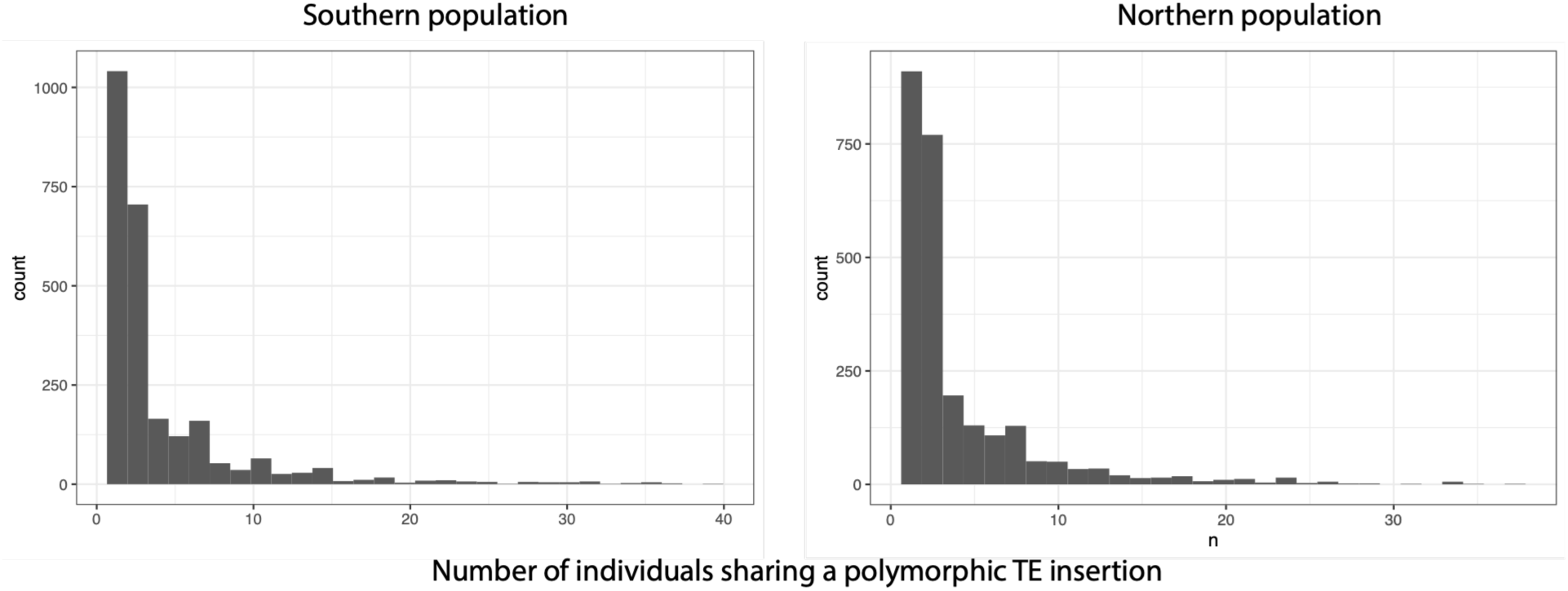
Number of individuals sharing polymorphic TE insertions in north and south *C. ariakensis* populations.

**Sup. Fig. 6:**
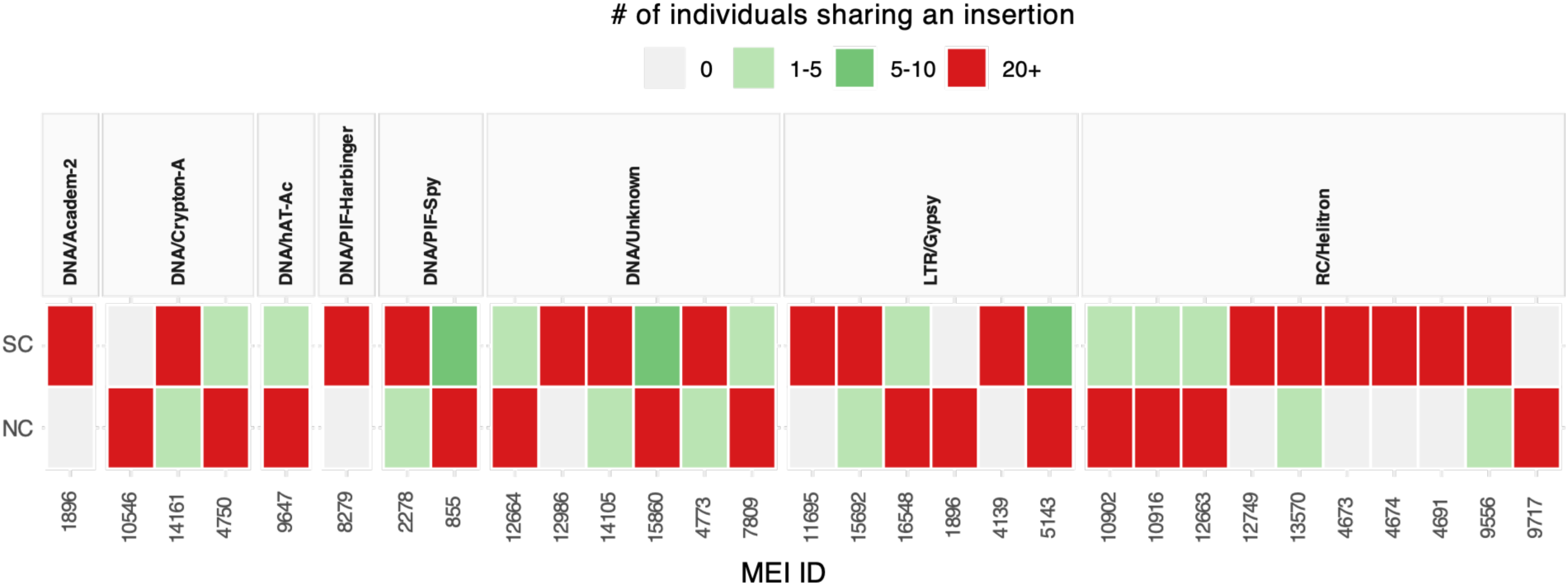
Number of individuals sharing the 30 MEIs exhibiting significant frequency differences between North (SC) and South (SC) *C. ariakensis* populations.

